# Stronger effects of heterozygosity on survival in harsher environments

**DOI:** 10.1101/177725

**Authors:** S Vincenzi, D Jesensek, JC Garza, AJ Crivelli

## Abstract

A stronger correlation between heterozygosity and fitness or its components (e.g., life-history traits such as survival, growth, morphology) is expected in harsher environments, but few studies have investigated whether the effects of heterozygosity on life-history traits vary with environmental conditions in natural populations. Here, the hypothesis that the effects of heterozygosity vary with environmental conditions was tested using six populations of marble trout *Salmo marmoratus* from Western Slovenia as a model system. Specifically, the tested hypotheses were: stronger effects of heterozygosity on survival in populations characterized by low average survival, no effects of heterozygosity on probability of surviving flash floods owing to their largely non-selective effects across traits, and stronger effects of heterozygosity on survival for fish born after floods than fish born before. A significant effect of heterozygosity on survival was found in populations characterized by low average survival. There were no effects of heterozygosity on probability of surviving flash floods, but in one population a positive correlation between heterozygosity and survival for fish born after the extreme events was found, probably because crowding in a small section of the stream caused more intense competition for resources.

## INTRODUCTION

Understanding within-population variation in components of fitness, and how they are affected by genetic variation, is a central topic in ecology and evolutionary biology (Chapman et al., 2009). The concept of individual quality has often been used to explain inter-individual heterogeneity in performance (Moyes et al., 2009; Wilson & Nussey, 2010; Bergeron et al., 2011). Heterogeneity in individual quality may result from complex interactions among genetic, maternal, environmental and demographic factors at different stages of life (Bergeron et al., 2011). In particular, the strength of the relationship between individual quality and either fitness or particular components of fitness (e.g., survival, growth) may depend on the quality of the environment. A stronger correlation between individual quality and fitness may be expected in harsher (e.g., food-poor, stronger competition for resources) environments (Reed et al., 2007), while in more favorable environments there may be enough resources available for all individuals to survive to sexual maturity and reproduce (Leung & Forbes, 1997). In addition, the effects of individual quality on survival may be greater during extreme events (e.g., cold spells, floods, fires) and in the post-extreme event environment, due to higher densities and stronger competition for resources (Keller et al., 1994; Jenouvrier et al., 2015).

Heterozygosity is often considered a proxy of individual quality and a positive correlation between heterozygosity and fitness-related traits has often been found in fishes (David, 1998; Pujolar et al., 2005). Multiple hypotheses on the proximate mechanisms of a positive heterozygosity-fitness correlation have been advanced (e.g., functional overdominance, inbreeding depression) and, as expected, the effect sizes are generally small (Chapman et al., 2009). Although investigations have been encouraged (Chapman et al., 2009), the hypothesis of whether the effects of heterozygosity on fitness components vary with environmental conditions has rarely been tested in natural populations of fish species. Here, this hypothesis was tested using as a model system six marble trout (*Salmo mamoratus* Cuvier 1829) populations living in Western Slovenia (Vincenzi et al., 2016). Average survival probabilities of marble trout differ greatly among populations and three of them have been affected recently by flash floods causing massive mortality (Vincenzi et al., 2016). Specifically, the hypotheses tested were: (a) positive effects of heterozygosity on survival in populations with lower average survival rates or when in competition with other species (i.e., harsher/poorer environment), (b) no effect of heterozygosity on surviving flash floods owing to their largely non-selective effects across traits, and (c) greater effects of heterozygosity for fish born after the occurrence of flash floods than born before, due to higher population density and stronger competition for resources after the flood.

## MATERIAL AND METHODS

### STUDY AREA AND SPECIES DESCRIPTION

Detailed descriptions of the biology and ecology of marble trout can be found in Vincenzi et al., (2016). The hypotheses on effects of heterozygosity on survival varying with environmental conditions were tested on the marble trout populations of Lower Idrijca [LIdri], Upper Idrijca [UIdri], Lipovesck [Lipo], Zadlascica [Zadla], Trebuscica [Trebu], and Zakojska [Zak]. In LIdri, marble trout live in sympatry with rainbow trout (*Oncorhynchus mykiss*) (Vincenzi et al., 2011). Three populations have been affected by flash floods, Lipo (flash floods in 2007 and 2009), Zadla (2007), and Zak (2007), leading to >55% decreases in survival with respect to sampling occasions with no floods (Vincenzi et al., 2016). In Zak, the only fish surviving the 2007 flood were living in a small stretch in the uppermost part of the stream, which was less affected by flows and debris (Vincenzi et al., 2017). Average survival probabilities (±se) in sampling occasions with no floods were: LIdri (0.38±0.01), UIdri (0.51±0.02), Lipo (0.45±0.06), Zadla (0.33±0.05), Trebu (0.5±0.02), Zak (0.49±0.01) (Vincenzi et al., 2016).

### SAMPLING

Populations were sampled either annually in June (Zak) or September (Zadla, Trebu) or biannually in both months (LIdri, UIdri, Lipo). Tagging started in different years for different populations: 1996 - Zak; 2004 - LIdri and UIdri; 2006 – Zadla, Lipo, Trebu. Sampling protocols are described in detail in Vincenzi et al. (2016). Fish were captured by electrofishing and fork length (*L*) and weight recorded to the nearest mm and g, respectively. If a captured fish had *L* > 110 mm, and had not been previously tagged or had lost a previously applied tag, it received a Carlin tag (Carlin, 1955) and age was determined by reading scales. The adipose fin was also removed from all fish captured for the first time, including those not tagged due to small size. Fish are aged as 0+ in the first calendar year of life, 1+ in the second year and so on. Sub-yearlings are smaller than 110 mm in June and September, so fish were tagged when at least aged 1+. We included data up to 2014 for each population.

### SNP DISCOVERY AND GENOTYPING

DNA was extracted from dried fin clips using the Dneasy 96 filter-based nucleic acid extraction system on a BioRobot 3000 (Qiagen, Inc.), following the manufacturer’s protocols. Extracted DNA was diluted 2:1 with distilled water and used for polymerase chain reaction (PCR) amplification of population-specific SNPs. SNPs were assayed with 96.96 genotyping IFC chips on an EP1 (EndPoint Reader 1) instrument (Fluidigm, Inc.), using the manufacturer’s recommended protocols. Genotypes were called using SNP Genotyping Analysis software (Fluidigm). Two people called all genotypes independently, and discrepancies in the scores were resolved either by consensus, by re-genotyping, or by deletion of that genotype from that data set. A proportion of individuals were sampled and genotyped more than once (e.g., in case of tag loss or when the fish was sampled when < 110 mm, and then later at tagging), as determined by observed identical genotypes and compatible age and length data, and one of the samples was excluded from the analyses. Matching genotypes of individuals with different tags is a method of “genetic tagging” that allows reconstructing the life histories of individuals after tag loss.

We used 118 SNPs for Lipo (Mean±sd minor allele frequency (MAF) of the SNPs: 0.23±0.15), 94 for Zak (0.28±0.13), 95 for LIdri (0.27±0.14), 95 for UIdri (0.26±0.14), 95 for Trebu (0.28±0.14), 95 for Zadla (0.30±0.11). Individual expected heterozygosity was calculated as the observed proportion of an individual’s loci that were heterozygous. The final dataset included 3416 marble trout total, from LIdri, n = 781; UIdri, 502; Lipo, 384; Zadla, 220; Trebu, 278 and Zak, 1241.

### SURVIVAL ANALYSIS

The goal was to estimate the effects of heterozygosity on variation in survival for each population while accounting for year of birth, and season and occurrence of flash floods, where applicable. The hypotheses are: (a) stronger effects of heterozygosity on survival in populations with lower survival probabilities (Zadla) or with marble trout living in sympatry with rainbow trout (LIdri), (b) no effect of heterozygosity on surviving flash floods and (c) a stronger effect of heterozygosity for fish born after, rather than before, the occurrence of flash floods in Zak and Lipo, due to higher juvenile production post-flood (Vincenzi et al., 2016, 2017) and consequent higher competition for resources (e.g., food, shelter). For (c), it was also predicted a stronger effect in Zak than in Lipo due to higher density/stronger competition of fish born after the flood in Zak (Vincenzi et al., 2017). For Zadla, the sample size was too small for testing hypothesis (c).

Two relevant probabilities can be estimated from a capture history matrix: *ϕ*, the probability of apparent survival, and *p*, the probability that an individual is captured given that it is alive (Thomson et al., 2009). The Cormack–Jolly–Seber (CJS) model was used as a starting point for the analyses (Thomson et al., 2009). Previous work has found no or minor effects of population density, water temperature, body size, or sex on survival in marble trout (Vincenzi et al., 2016).

For each population, the best recapture model as reported in Vincenzi et al. (2016) was used. A seasonal effect (*Season*) was modeled as a simplification of full time variation, dividing each year in two periods: June to September (*Summer*) and September to June (*Winter*). Since the length of the two intervals (*Summer* and *Winter*) was different, the probability of survival was estimated on a common annual scale, including *Season* as a potential predictor of probability of capture and survival in all populations that have been sampled twice a year. *Flood* (0 for no flood occurring during the sampling interval and 1 otherwise) was included as a potential predictor of probability of survival for Lipo, Zak, Zadla.

Fish with fewer than 50 SNPs (∼half of total SNPs across populations) genotyped were excluded from the analysis. Heterozygosity was not standardized within populations since values on their natural scale are more easily interpretable and the ranges of individual heterozygosities were very similar among population. Previous studies on marble trout have shown that a large fraction of the variability in survival is explained by year of birth, season, and the occurrence of floods (Vincenzi et al., 2016). Therefore, models tested included a year-of-birth component (*year-class*, *pre-flood or post-flood*, constant), a time component (*Season*, *Flood*, or constant) and a heterozygosity component (linear, cubic splines, or constant). Year-class is the year of birth of a fish, and *pre-flood/post-flood* is a categorical variable identifying fish born before or after the flash floods in Lipo and Zak. Since just a few fish were born in Lipo between the flash floods of 2007 and 2009 (Vincenzi et al., 2017), fish born in 2008 and later in Lipo were *post-flood*. We did not include sampling occasion in our model of survival, in order to avoid it masking the role of other determinants of survival of expected much smaller effect size (such as heterozygosity).

For each model, we tested additive and multiplicative interactions among predictors (see R code for full list of models). Models were considered to provide the same fit to data when ΔAIC between them was < 2 and, in the case of multiple models with ΔAIC < 2, the model with the fewest parameters was selected as the most supported (Burnham & Anderson, 2002). We did not use null hypothesis significance testing for model and variable selection, and heterozygosity was thus assumed to explain variation in survival (relative to models not including heterozygosity) when it was included in the most supported model. We carried out the analysis of survival using the package *marked* (Laake et al., 2013) for R (R Development Core Team, 2014).

Data and code are available at https://github.com/simonevincenzi/Heter

## RESULTS

Models including heterozygosity as a predictor of survival were strongly supported in LIdri, Zadla, Trebu and Zak (Table 1). In Zak, the effects of heterozygosity varied with year of birth (i.e., before or after the flash flood of 2007; Table I). For these populations, the relationship between heterozygosity and survival was positive (Fig. 1), although in LIdri a cubic spline mostly predicted very low survival probabilities for fish with low heterozygosity (Fig. 1d). No effect of heterozygosity on fish survival was observed in Lipo, Zadla or Zak for the sampling occasion in which flash floods occurred (Fig. 2), but a positive effect of heterozygosity on fish survival was observed in Zak after the flash flood of 2007 (Table 1, Fig. 1a).

**Fig. 1.**
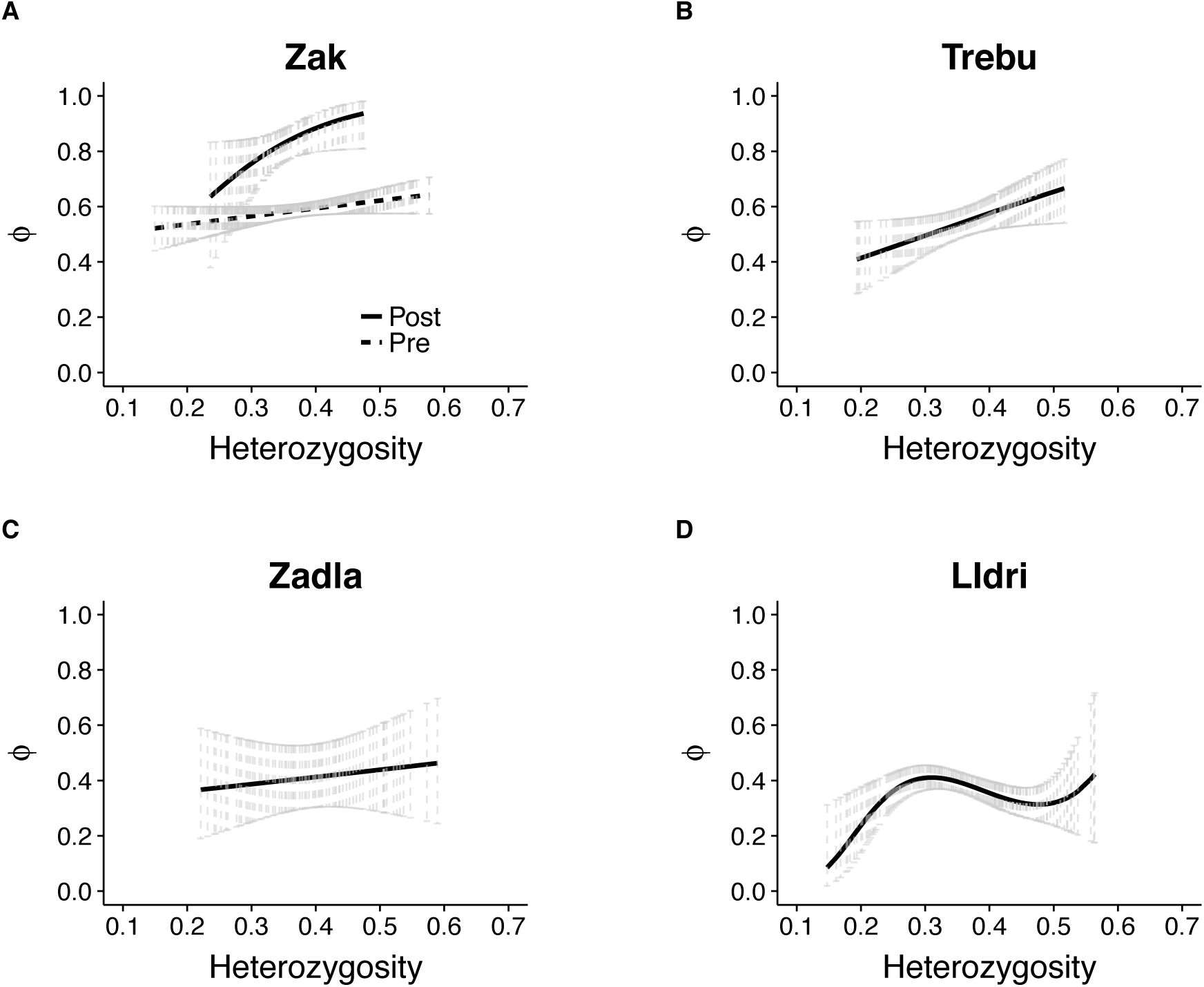
Relationship between heterozygosity and apparent survival *ϕ* in the four marble trout populations for which models including heterozygosity were strongly supported. Vertical lines are 95% CIs. For Zak, Pre: fish born before the 2007 flash flood, Post: fish born after the 2007 flash flood. For both Zak and Zadla, survival probabilities are for sampling occasions with no extreme event.

**Fig. 2.**
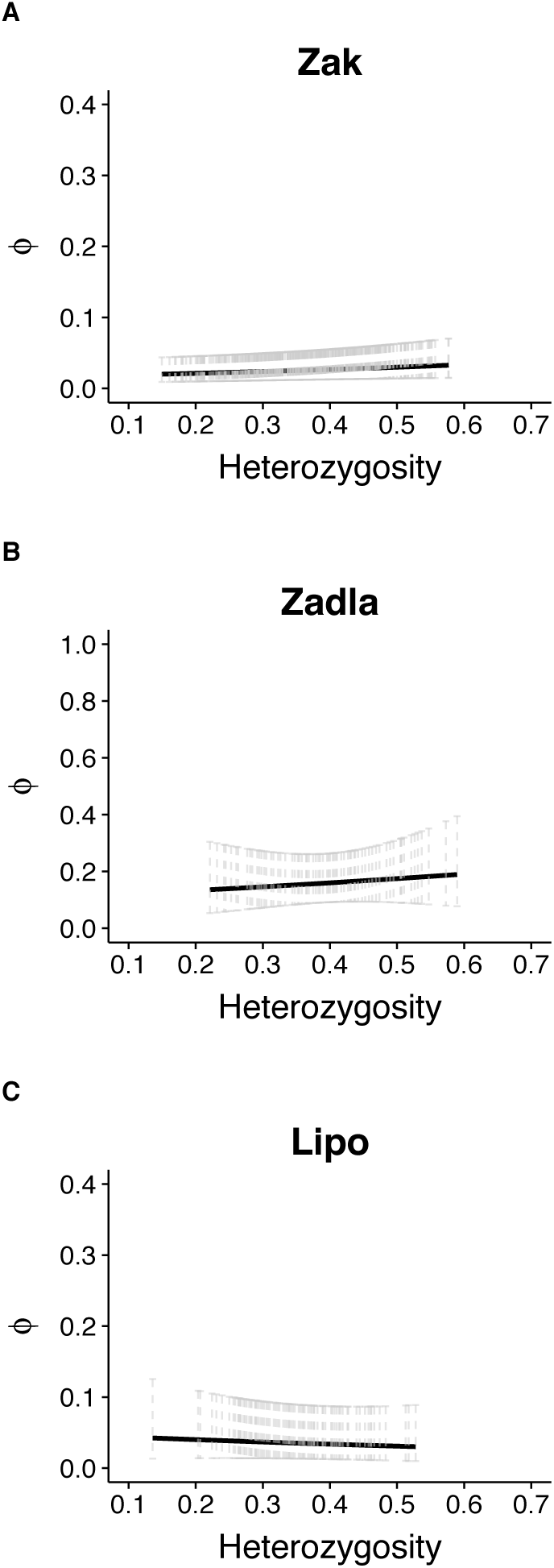
Relationship between heterozygosity and probability of surviving a flash flood in the populations of Zakojska, Zadlascica, and Lipovscek (see online code for model ranking and parameter estimates)

**Table 1.** Population-specific models of apparent survival *ϕ* within ΔAIC < 2 from the best model. The symbol * denotes interaction between predictors. *time* = interval between two consecutive sampling occasions (only used for recapture models). *Season* = categorical variable for *Summer* (June to September) and *Winter* (September to June). *heter* = expected individual heterozygosity. *Pre.Post* = categorical variable for fish born before (*Pre*) or after (*Post*) flash floods. *bs* = b-splines. *npar* = number of parameters of the survival model.

| Model | *npar* | $\Delta AIC$ | *weight* |
| --- | --- | --- | --- |
| **Zak** |  |  |  |
| $\phi(heter * Pre.Post + Flood)p(Age)$ | 7 | 0.00 | 0.51 |
| $\phi(heter + Pre.Post + Flood)p(Age)$ | 6 | 0.95 | 0.32 |
| **Trebu** |  |  |  |
| $\phi(heter) p(time)$ | 10 | 0.00 | 0.54 |
| $\phi(bs(heter) p(time))$ | 12 | 1.23 | 0.29 |
| **Zadla** |  |  |  |
| $\phi(heter + Flood)p(const)$ | 4 | 0.00 | 0.80 |
| **Lidri** |  |  |  |
| $\phi(bs(heter)) p(bs(Age))$ | 8 | 0.00 | 0.43 |
| $\phi(bs(heter) + Season) p(bs(Age))$ | 9 | 1.15 | 0.24 |
| **UI dri** |  |  |  |
| $\phi(const) p(bs(Age))$ | 5 | 0.00 | 0.30 |
| $\phi(bs(heter)) p(bs(Age))$ | 8 | 1.16 | 0.17 |
| $\phi(heter)p(bs(Age))$ | 6 | 1.42 | 0.15 |
| $\phi(Season)p(bs(Age))$ | 6 | 1.44 | 0.15 |
| **Lipo** |  |  |  |
| $\phi(Pre.Post + Flood)p(const)$ | 26 | 0.00 | 0.98 |

## DISCUSSION

The results of this study confirmed the hypothesis of stronger effects of heterozygosity on survival in poorer environments, with the exception of the population of Trebuscica, for which the most-supported model predicted a positive effect of heterozygosity on survival, although average survival was the second highest among the 6 marble trout populations.

The consequences of extreme events on trait selection are still poorly understood; in some cases, all phenotypes are affected equally by an extreme event, while in others, some phenotypes may respond better than others to the unusually harsh conditions, which thus act as an intense selection episode. For instance, song sparrows (*Melospiza melodia*) that survived a severe population bottleneck caused by a harsh winter were a non-random subset of the pre-crash population with respect to inbreeding, indicating that natural selection favored outbred individuals (Keller et al., 1994). However, flash floods and debris flows such as those that are occurring with increasing frequency in Western Slovenia are unlikely to directly select for particular phenotypes through differential survival, due to their catastrophic, unpredictable effects (Vincenzi et al., 2017). Consistent with this, it was not found that more heterozygous fish had higher chances of surviving flash floods.

In Zakojska, a strong, positive relationship between heterozygosity and survival was found for fish born after the 2007 flash flood. In contrast, no effects of heterozygosity on survival was found for fish born after the flash floods in Lipovscek. High average survival for fish born after the flash flood may indicate particularly favorable environmental conditions; however, high survival may also be due to a combination of lower cannibalism and fewer older fish competing for space, while intra-cohort competition for food and space may still be intense. The different relationships found in the two populations are consistent with the hypothesis that a stronger correlation between individual quality and fitness is expected in harsher environments. In Zakojska, crowding in a small section of the stream after the 2007 flash flood caused more intense competition and may have contributed to stronger selection for heterozygosity than prior to the flood.

We thank the employees and members of the Tolmin Angling Association (Slovenia) for carrying out fieldwork since 1993. We also thank E. Campbell, C. Columbus and E. Correa for assistance with genetic data generation.

